# Bacteriophage Φ21’s receptor-binding protein evolves new functions through destabilizing mutations that generate non-genetic phenotypic heterogeneity

**DOI:** 10.1101/2024.04.15.589635

**Authors:** Krista R. Gerbino, Joshua M. Borin, Sarah M. Ardell, Justin J. Lee, Kevin D. Corbett, Justin R. Meyer

**Affiliations:** School of Biological Sciences, University of California San Diego, La Jolla, California, USA; Department of Cellular and Molecular Medicine, University of California San Diego, La Jolla, California, United States of America

## Abstract

How viruses evolve to expand their host range is a major question with implications for predicting the next pandemic. Gain-of-function experiments have revealed that host-range expansions can occur through relatively few mutations in viral receptor-binding proteins, and the search for molecular mechanisms that explain such expansions are underway. Previous research on expansions of receptor use in bacteriophage λ has shown that mutations that destabilize λ’s receptor-binding protein cause the receptor-binding protein to fold into new conformations that can utilize novel receptors but have weakened thermostability. These observations led us to hypothesize that other viruses may take similar paths to expand their host range. Here, we find support for our hypothesis by studying another virus, bacteriophage 21 (Φ21), which evolves to use two new host receptors within two weeks of laboratory evolution. By measuring the thermodynamic stability of Φ21 and its descendants, we show that, as Φ21 evolves to use new receptors and expands its host range, it becomes less stable and produces viral particles that are genetically identical but vary in their thermostabilities. Next, we show that this non-genetic heterogeneity between particles is directly associated with receptor use innovation, as phage particles with more derived receptor use capabilities are more unstable and decay faster. Lastly, by manipulating the expression of protein chaperones during Φ21 infection, we demonstrate that heterogeneity in thermostability and receptor use breadth of phage particles is directly related to the folding of phage proteins into different conformers. Altogether, our results provide support for the hypothesis that viruses can evolve new receptor-use tropisms through mutations that destabilize the receptor-binding protein and produce multiple protein conformers.

## Introduction

As we enter the fourth year of a global pandemic, many researchers have turned their attention to understanding the risk factors and processes involved in potentiating pandemics (Franch-Pardo et al., 2020). A critical step in the initiation of a pandemic occurs when non-human viruses gain the capacity to recognize and bind to human cells (Ito et al., 1998). Often, this occurs when viruses acquire specific mutations in their receptor-binding protein (RBP), which allows them to attach to molecules on the surface of new host cells (Longdon et al., 2014). Since the beginning of the SARS-CoV-2 pandemic, there has been renewed interest in 1) understanding the types of evolutionary and molecular processes that lead to changes in RBP tropism and 2) determining whether there are general processes by which these molecular innovations evolve.

Numerous research groups have investigated how RBPs evolve to use new receptors. Many studies have reported lists of specific mutations (Curiel, 2006; Grimm et al., 2008; Meyer et al., 2012) or other mutational pathways, like gene duplications (Tétart et al., 1996; Scholl et al., 2001), which led to gain-of-function in their study system. Yet, generalizable principles or mechanisms that explain how RBPs evolve new functions are rarely reported. One exception is a series of studies of the evolution and host range expansion of bacteriophage λ. When λ is cocultured in the laboratory with its host, *Escherichia coli*, λ rapidly and repeatedly evolves to use a new receptor, OmpF, in addition to its original receptor, LamB, by acquiring four or more mutations in its RBP called J (Meyer et al., 2012). Before gaining the ability to infect using OmpF, the earliest J mutations that λ evolves increase its adsorption to its native receptor, LamB (Burmeister et al., 2016). These mutations also pleiotropically destabilize the J protein, resulting in new thermodynamic properties that cause a single peptide sequence to fold into multiple distinct protein conformations, termed non-genetic phenotypic heterogeneity (Petrie et al., 2018; Strobel et al., 2022). After the RBP has been destabilized, λ evolves additional mutations along the RBP-receptor interface that confer the ability to use the new OmpF receptor (Strobel et al., 2022). Altogether, these observations led us to hypothesize that viral RBPs can gain new receptor tropisms by evolving mutations that destabilize the protein and produce a variety of protein conformers that can interact with new cellular features and explore new functions.

This hypothesis was inspired by the 70-year-old hypothesis from C. H. Waddington, a fruit fly evolutionary biologist who suggested that phenotypic novelty could first arise from mechanisms that induce phenotypic heterogeneity (e.g., phenotypic plasticity or other non-genetic mechanisms), which are then selected and canalized into new functions (Waddington, 1942). Recently, this idea has gained prominence and appears to be responsible for evolutionary innovations in several systems, including protein evolution (Van Buskirk and Steiner, 2009; Levis and Pfennig, 2019; Sakuma et al., 2023).

However, many questions remain about how the evolution of different protein conformers allow viral RBPs to acquire new tropisms. Here we address two: 1) does this mechanism apply to the evolution of other viruses besides λ, and 2) does the non-genetic heterogeneity arise during protein folding? Stochasticity in protein folding was implicated in previous work on λ because mutations in the RBP produced phenotypically heterogenous particles; however, the role of protein folding was not directly tested.

We tackle the first question by studying phage Φ21, which was recently reported to evolve to use two new receptors, OmpC and OmpF, in addition to its native receptor, LamB, when cocultured with *E. coli* hosts (Borin et al., 2023). Φ21 is related to phage λ, but only shares 94% nucleotide identity with λ’s RBP, and therefore represents a second test of this hypothesis (Ye et al., 2006).

We tested whether Φ21 evolved non-genetic phenotypic heterogeneity as it gained new receptor-use functionality by first focusing on a single Φ21 isolate (ΦD9) that was isolated on the ninth day of the previous experiment and was the earliest strain isolated that could use all three receptors. ΦD9 evolved four mutations in its RBP, another in a separate tail fiber protein, and one in a non-tail structural gene (Borin et al., 2023). We started by assessing whether an isogenic stock of this phage showed multiphasic decay, which was the first phenomenon observed in the evolved isolates of phage λ that led to this line of research (Petrie et al., 2018). If a protein folds into multiple conformations, there is a chance that the different conformations vary in their intrinsic thermodynamic properties and spontaneously decay at different rates. Additionally, if the decay of the RBP causes loss of particle infectivity, as observed with λ, then phenotypic heterogeneity can be detected by enumerating the loss of viable phage (measured as plaque forming units) over time in a phage lysate deprived of hosts, to test whether the phage’s decay has multiple phases instead of a single exponential decay rate.

We found evidence that ΦD9 exhibits a biphasic decay pattern and then investigated whether heterogeneity in particle decay rates is linked to heterogeneity in receptor use by measuring whether the population lost the ability to use different receptors at different rates. If all particles have the same receptor affinity, then the loss should be uniform; however, we found that the most unstable particles use OmpF, then OmpC, and the most stable particles can only infect through LamB.

Next, we investigated the second question, about whether there is a link between receptor-use heterogeneity and the process of protein folding. To do this, we manipulated protein folding within host cells by inducing heat-shock protein-folding chaperones and tested whether the presence of chaperones impacts the relative frequencies of Φ21 particles that use the three different receptors. We predicted that more chaperones would result in an increased frequency of proteins folded into the stable protein conformation that can only use LamB. In line with our prediction, we detected a statistically significant increase in stable LamB-using particles and a measurable decline in unstable OmpF- and OmpC-using particles.

Lastly, we tested how decay rates and multiphasic decay patterns change throughout Φ21’s evolution. After Φ21’s initial expansion to use 3 host receptors, phage receptor-use contracts along two different evolving lineages; one lineage evolves into dual-receptor phages that use LamB and OmpC (and then the lineage goes extinct) (Borin et al., 2023). The other lineage loses OmpC use and then LamB use, and eventually specialized on the OmpF receptor. We studied 7 phage isolates that span this diversity of receptor-use tropisms and found that although decay rates vary substantially, biphasic decay patterns were maintained throughout Φ21’s evolution.

## Methods

### Strains

All phage isolates used in this study are descendants of a strictly lytic version of Φ21 (GenBank: OL657228; Feiss et al., 2022) and were first reported in Borin et al., 2023. Relevant information about the phages used in this study are provided in Table 1. For bacterial hosts, *E. coli* K-12 strain BW25113 (WT), and related gene knockout mutants, were used for culturing and plating phages (Baba et al., 2006; gene double- and triple-receptor knockout strains described in Borin et al., 2023). For experiments where we manipulated host protein-folding chaperones, we used ASKA strain JW3426 (National BioResource Project (NIG, Japan)), which has a full-frame knockout of gene *rpoH* (sigma32) and a complemented version of the gene provided on the pCA24N plasmid under the control of an IPTG-inducible T5-lac promoter (Kitagawa et al., 2006).

**Table 1.**
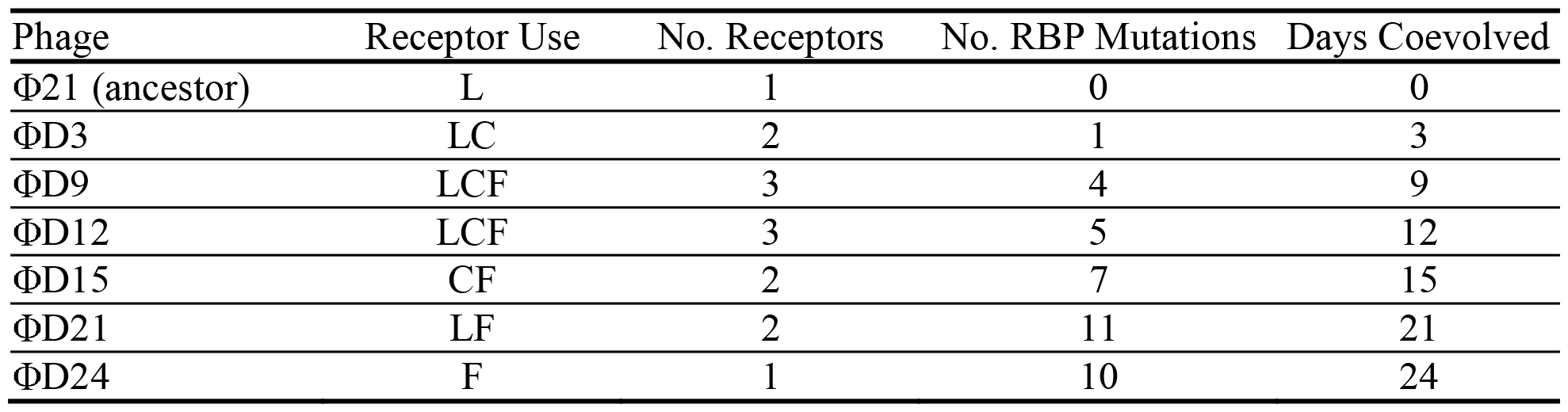
Relevant characteristics of phage genotypes included in this study. All phages are descendants of the Φ21 ancestor and were isolated from the same population. Additional information on these phages is reported in Borin et al., 2023.

### Phage ΦD9 decay experiment

To investigate whether Φ21 evolved phenotypic heterogeneity as it expanded the range of receptors it could use, we compared the decay patterns of the Φ21 ancestor with ΦD9. Phage decay was measured by first generating replicate isogenic phage lysates. Scrapes (~2 uL) from a frozen preserved isogenic phage stock were inoculated into replicate tubes containing 4 mL Tris-LB and ~10^7^ host cells (recipe in Borin et al., 2023). ΦD9 was grown with ΔLamB ΔOmpC host cells in order to maintain its ability to use the OmpF receptor. The Φ21 ancestor was grown with ΔOmpC cells (KEIO strain JW2203) to prevent its evolution to use the OmpC receptor. After incubating co-cultures for 4 h, shaking at 37°C, phages were extracted using 0.2 μm filters.

Phage lysates were then studied immediately to limit decay of phages. Phages were first diluted in Tris-LB and ~3×10^5^ particles were aliquoted into replicate glass tubes and incubated at 37°C without shaking to reduce evaporation. Initially and every day for 7 days, 100 μL was removed from each tube and inoculated into molten (~55°C) 0.7% w/w LB soft agar (recipe in Borin et al., 2023) infused with ~10^8^ WT cells, swirled onto LB agar plates, and incubated overnight at 37°C. After incubation, plaques were enumerated. We used WT because it expresses all 3 relevant host receptors and provides an estimate of all phage particles. The decay experiment was performed on a total of 12 replicates conducted in two separate batches. To compare the overall decay rate of the Φ21 ancestor and the evolved ΦD9 phage, we computed the exponential decay rate of each replicate between day 0 and 7 (Equation 1, below) and compared the rates using a T-test, implemented in R (version 4.1.1; R Core Team, 2021).

### Population decay model fitting

To study whether decay patterns of phages were suggestive of a single, monomorphic population of phage particles or a heterogeneous population comprised of particles with different decay rates, we used linear regression and model fitting (Fig. S1, Table S1). We fit models that were representative of populations containing a single decay rate (Eq. 1; monophasic), heterogeneous populations containing two (Eq. 2; biphasic) or three (Eq. 3; triphasic) discrete decay rates, and a model where decay rate changed continuously over time (Eq. 4, derived from Eq. 5) (Petrie et al., 2018). After fitting each model to the observed data, we calculated the Akaike Information Criterion (AIC) and log-likelihood to select the most supported model for our data (e.g., lowest AIC, a composite of fit and model complexity; significant P-value in model comparison, significant increase in fit).

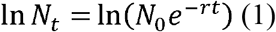

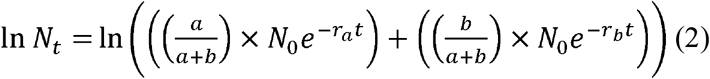

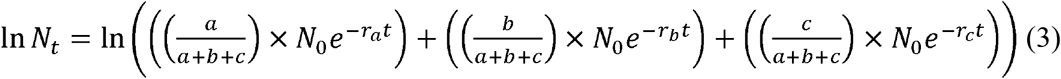

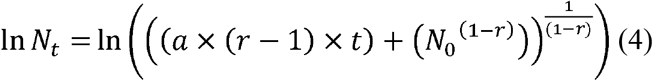

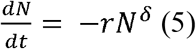

In order to test whether the decay of evolved ΦD9 is linked to subpopulations of phage particles with different receptor-use tropisms, we repeated the 7-day decay experiment described above except that at each timepoint we plated ΦD9 onto a suite of double-receptor knockout hosts (ΔOmpC ΔOmpF, ΔLamB ΔOmpF, and ΔLamB ΔOmpC), to enumerate the particles that remained viable and could infect through each different receptor. We then calculated the proportion of phage particles that could use each receptor by calculating the efficiency of plaquing (EOP) by dividing the density of phage particles that could use a given receptor by the total density of phage particles (determined by enumerating the phage lysate on WT cells which present all three receptors). To compare the decay rate of each subpopulation that could infect a different receptor, we calculated initial exponential decay rates (between 0 and 48 h) and then analyzed these rates using ANOVA and Tukey’s HSD tests, implemented in R.

### Protein folding and host receptor shift experiment

To test whether there is a link between the heterogeneity of phage receptor use and phage protein folding, we modified the expression of host heat-shock chaperone proteins during infection to test if this would alter phage protein folding and produce phages with different receptor-use preferences. To do this, we grew *E. coli* cultures by inoculating a colony of JW3426 grown on an agar plate (LB agar with 0.1 M chloramphenicol and 0.8 M IPTG) into tubes containing 4 mL LB supplemented with the same concentrations of chloramphenicol and IPTG, and incubated the cultures overnight, shaking at 30°C. The next day, cells were washed 4 times by pelleting cells via centrifugation (21,130 x *g* for 2 min) and resuspending them in Tris-LB. After washing, ~10^7^ cells were inoculated into 4 mL LB with chloramphenicol but without IPTG and incubated overnight, shaking at 37°C. After incubation, cells were washed again, as described above, to remove any remaining IPTG. Finally, ~10^7^ cells were inoculated into 12 replicate tubes containing 4 mL Tris-LB supplemented with 0.1M chloramphenicol and ~10^4^ particles of ΦD9. Half of the replicates were also supplemented with 0.8 M IPTG and half were not (n=6 replicates in each treatment). Cultures were incubated at 37°C, allowing phages to replicate on each of the hosts in each treatment, and after 2 hours phages were extracted using 0.2 μm filters, serially diluted, spotted on LB agar plates infused with WT or our collection of double receptor knockout hosts, and incubated overnight at 37°C. The next day, we enumerated plaques and calculated the EOP. The entire experiment was conducted twice, and no day effects were observed (p>0.05). To determine whether there were differences in host receptor preferences between phages grown on host cells with or without *rpoH* expressed, we compared the EOP between treatments on each receptor using T-tests, implemented in R.

### Decay of evolved phages with diverse receptor-use tropisms

During coevolution with its *E. coli* host, Φ21 diversifies into phage strains with different receptor use capabilities (conferred by the acquisition of different numbers of mutations in the RBP, called J). To investigate whether and when receptor use expansion and contraction was associated with protein destabilization and the creation of phenotypic heterogeneity, we measured the decay of representative phage isolates with different receptor use capabilities (isolated from the earliest day on which that receptor use type was observed). Decay experiments were conducted as described above, except with the following changes: Phages ΦD3, ΦD9, ΦD12, ΦD15, ΦD21, ΦD24 were grown up from frozen preserved stocks with WT, ΔLamB ΔOmpC, WT, ΔOmpC, ΔLamB, or ΔLamB ΔOmpC hosts, respectively. We chose hosts that produced a high density of phage particles while also maintaining the receptor use range of the phage isolate. Experiments were conducted for 3 days, which encompassed >93% of the decay that was observed in previous experiments on ΦD9. To test whether protein stability was related to receptor use breadth, we fit linear models comparing the initial exponential decay rate (0–24 h) and the number of receptors each phage strain could use. Similarly, we compared phage stability with the number of evolved mutations in the phage’s RBP, as well as against the number of days the phage isolate had evolved.

### RBP structure predictions

To study how a key RBP mutation (I1025T) affects Φ21 stability, we used AlphaFold 2 to predict the structure of the C-terminal 319 amino acids of Φ21 RBP that includes the central straight fiber domain (CSF) and receptor-binding domain (RBD) of the protein, as a homotrimer. We used the publicly available ColabFold-mmseqs2 implementation (Mirdita et al., 2022) of AlphaFold-multimer (Jumper et al., 2021; Evans et al., 2022). We used PyMOL (Schrodinger Inc.) and Chimera (Pettersen et al. 2004) to pinpoint where the mutation occurs and to visualize how it could affect protein stability.

## Results

### Decay dynamics of the Φ21 ancestor and triple receptor-using ΦD9

We found strikingly different decay patterns for the Φ21 ancestor and ΦD9, which are separated by only four RBP mutations. ΦD9 decayed significantly faster than its ancestor (p<0.0001, T-test comparing initial decay rates; Fig. 1A). In addition, we found that the ancestor appears to be monomorphic because it decays at a single exponential rate, whereas ΦD9 was best fit by a biphasic exponential decay model, suggesting that the isogenic phage lysate was comprised of at least two phage subpopulations with different decay rates (Fig. 1A; Supplementary Material, Figure S1, Table S1).

**Figure 1.**
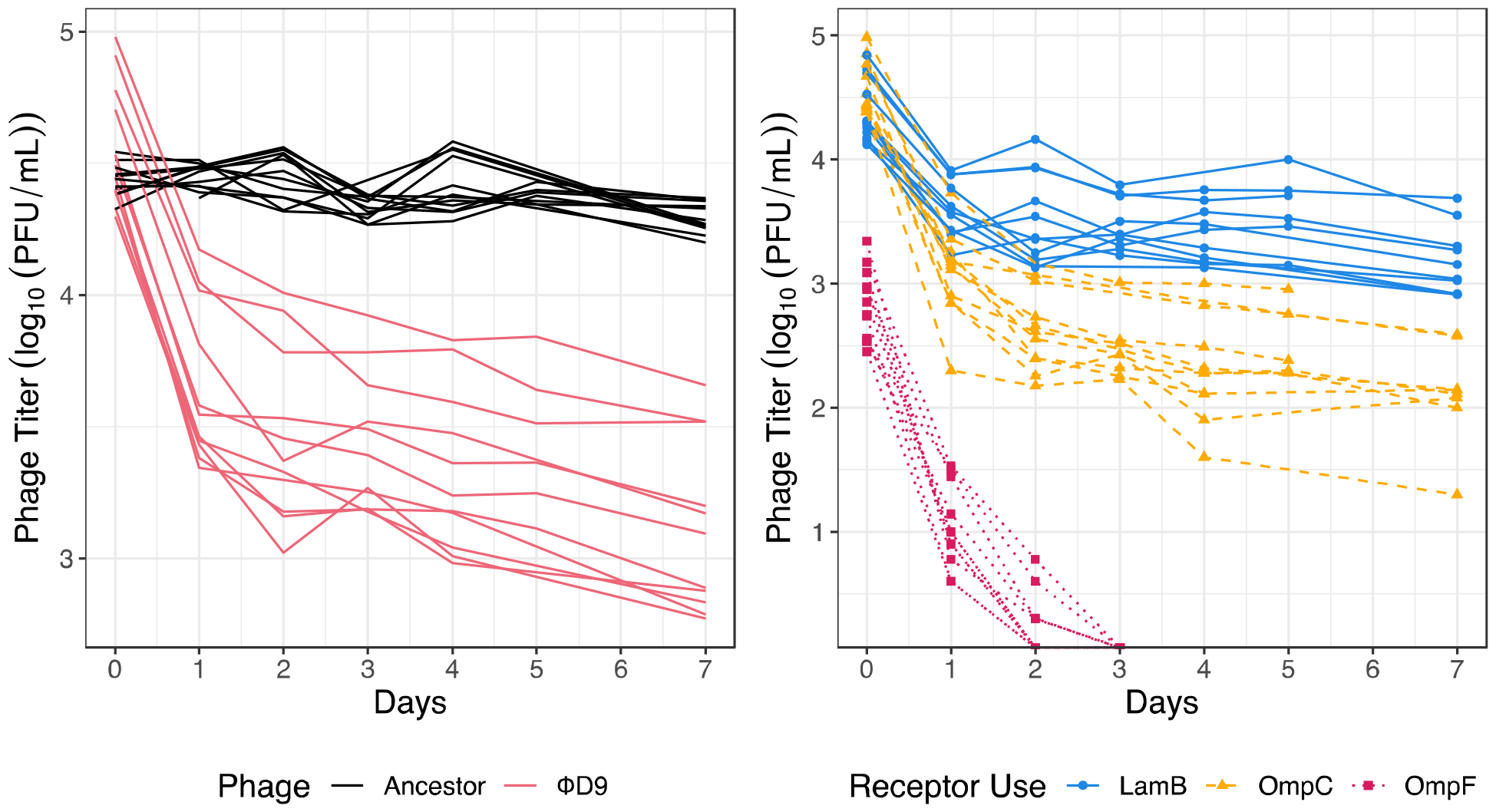
Phage Φ21 evolves multiphasic decay concomitant with host receptor use expansion. Multiphasic decay is linked to subpopulations of phage particles with different host receptor tropisms. Panel A: Titer of Φ21 ancestor (black) and evolved triple-receptor using phage ΦD9 (pink) decaying over 7 days. A monophasic decay model was the best fit for the ancestor and a biphasic decay model was the best fit for ΦD9 (Table S1). Panel B: Titer of subpopulations of phage particles that can infect using different host receptors. Lines represent individual replicates. Phage particle subpopulations with different receptor-use tropisms had significantly different initial decay rates (ANOVA, p<0.0001; Tukey’s HSD, p<0.01).

**Figure 2.**
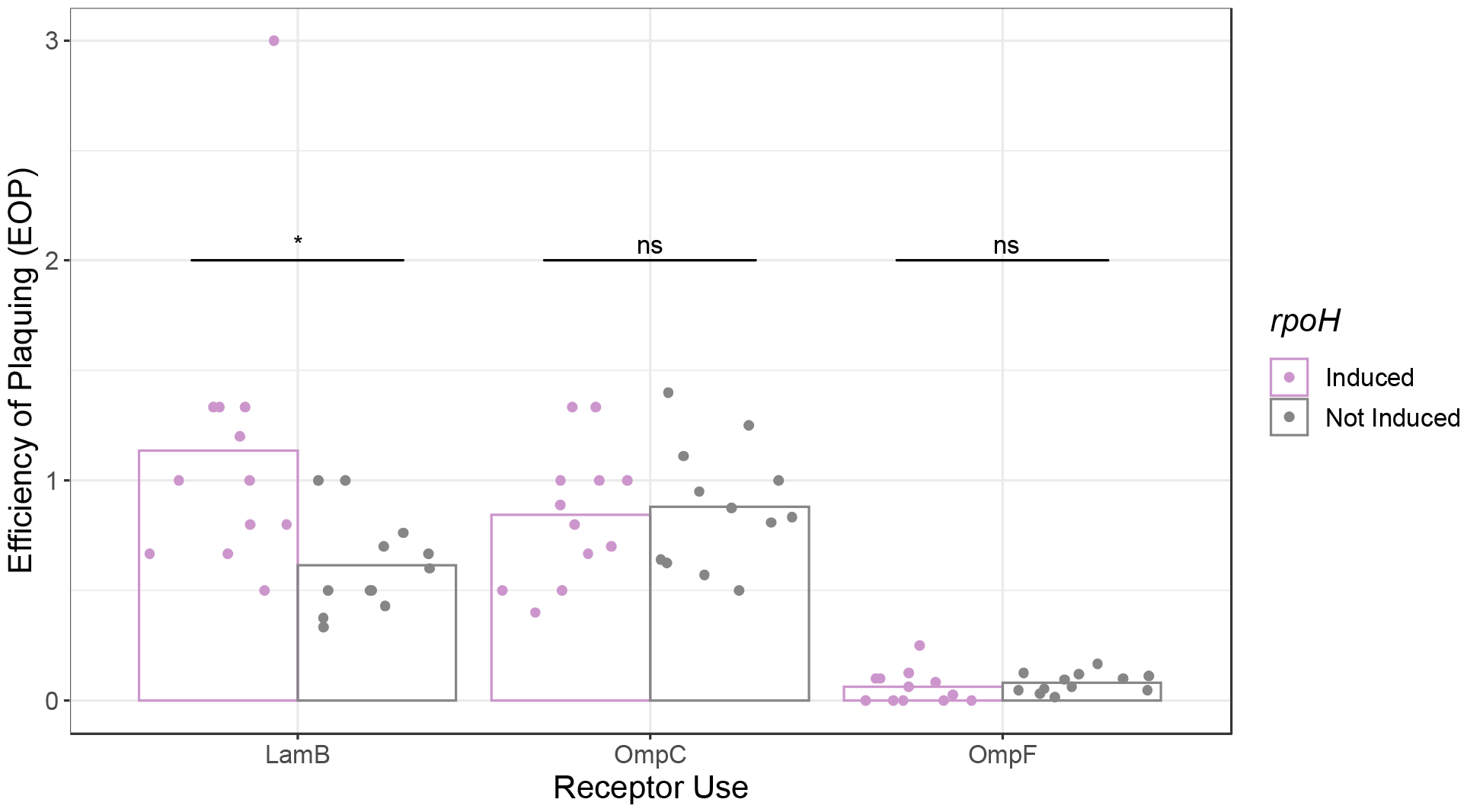
Increased expression of chaperone proteins (*rpoH* induction) shifts particle formation towards stable, LamB-using variants. Efficiency of plaquing (EOP) was calculated as the relative titer of phages on each receptor (i.e., single-receptor hosts) compared to on all receptors (i.e., WT). Significant differences in EOP treatments for each receptor are indicated above bars (*, p<0.05; ns, not significant; T-test; n=12 replicates per treatment).

In support of our hypothesis that the observed heterogeneity in particle decay rates underlies functional heterogeneity in receptor use, we found that subpopulations with different receptor preferences decayed at different rates. Particles that can use the native receptor (LamB) were the most stable, followed by particles that use the second receptor (OmpC), and the least stable were particles that use the third receptor, OmpF (Fig. 1B; ANOVA, p<0.0001; Tukey’s HSD, p<0.01). Interestingly, this reflects the order in which the new receptor-use functions evolved. The three significantly different decay rates measured from isogenic stocks of ΦD9 suggest that the isogenic ΦD9 culture has at least three different protein conformations with three different decay rates (Fig 1B). However, we were unable to detect triphasic decay with our curve-fitting analysis. We believe this was due to technical limitations, since the OmpF^+^ phage particles comprise only ~1% of the total particles and were only detected for two days, limiting their ability to drive a signal when intermixed within the full population of phage particles (Fig. 1B). Regardless of these technical limitations, the different thermostabilities associated with different receptor-using phage particles supports that Φ21 evolved to use novel receptors by producing a multimorphic population of particles in accordance with our hypothesis. This result provides a second non-λ example of RBP gain of function achieved through destabilizing mutations that cause non-genetic phenotypic heterogeneity.

### Link between protein folding and non-genetic phenotypic heterogeneity

We hypothesized that heterogeneity in phage particles arises due to stochasticity in the folding of RBPs in the host cell. To test our hypothesis, we altered the presence of protein-folding chaperones in the host during phage infection by manipulating expression of *rpoH*. In support of our hypothesis, we found that when *rpoH* was induced, the EOP for stable particles that use the LamB receptor was significantly higher than in cells without *rpoH* induced (T-test, p=0.02). In addition, the EOP for less stable particles that use OmpC and OmpF receptors were lower, although this difference was not statistically significant (p=0.76 for OmpC, p=0.46 for OmpF). One *rpoH*-induced datapoint showed a considerably higher EOP on LamB that other replicates, calling into question whether the difference observed was driven by an outlier. However, removing this point strengthens the statistical significance of our results (T-test, p=0.005). By directly perturbing the link between protein folding and the production of phenotypic heterogeneity, these experiments provide mechanistic support for our hypothesis that heterogeneity in RBP folding results in the formation of a range of conformers with different receptor-use functionalities.

### Stability changes during Φ21 coevolution

Thus far, we have focused on comparing the Φ21 ancestor and a derived variant (ΦD9) that evolved to use three receptors. For the rest of the study, we explore whether other evolved variants demonstrate similar decay patterns. We studied five new phages isolated from different timepoints or with unique receptor usages (Table 1). We measured the decay of these new evolved phages at three timepoints (0, 24, and 72 h) by enumerating the total phage density by plating on WT cells. These timepoints were sufficient to characterize the overall differences in decay rates, as well as to test whether strains exhibited multiphasic decay by comparing decay rates calculated from 0-24 h and 24-72 h. To ensure that these results were comparable with previous experiments, we also included ΦD9. We found that all evolved phages were less stable than the ancestor and they all show patterns that suggest biphasic decay (Fig. 3; paired T-test of difference in decay rate from 0-24 h and 24-72 h. ΦD3, p<0.001; ΦD9, p<0.0001; ΦD12, p<0.0001; ΦD15, p<0.0001; ΦD21, p<0.0001; and ΦD24, p<0.005).

**Figure 3.**
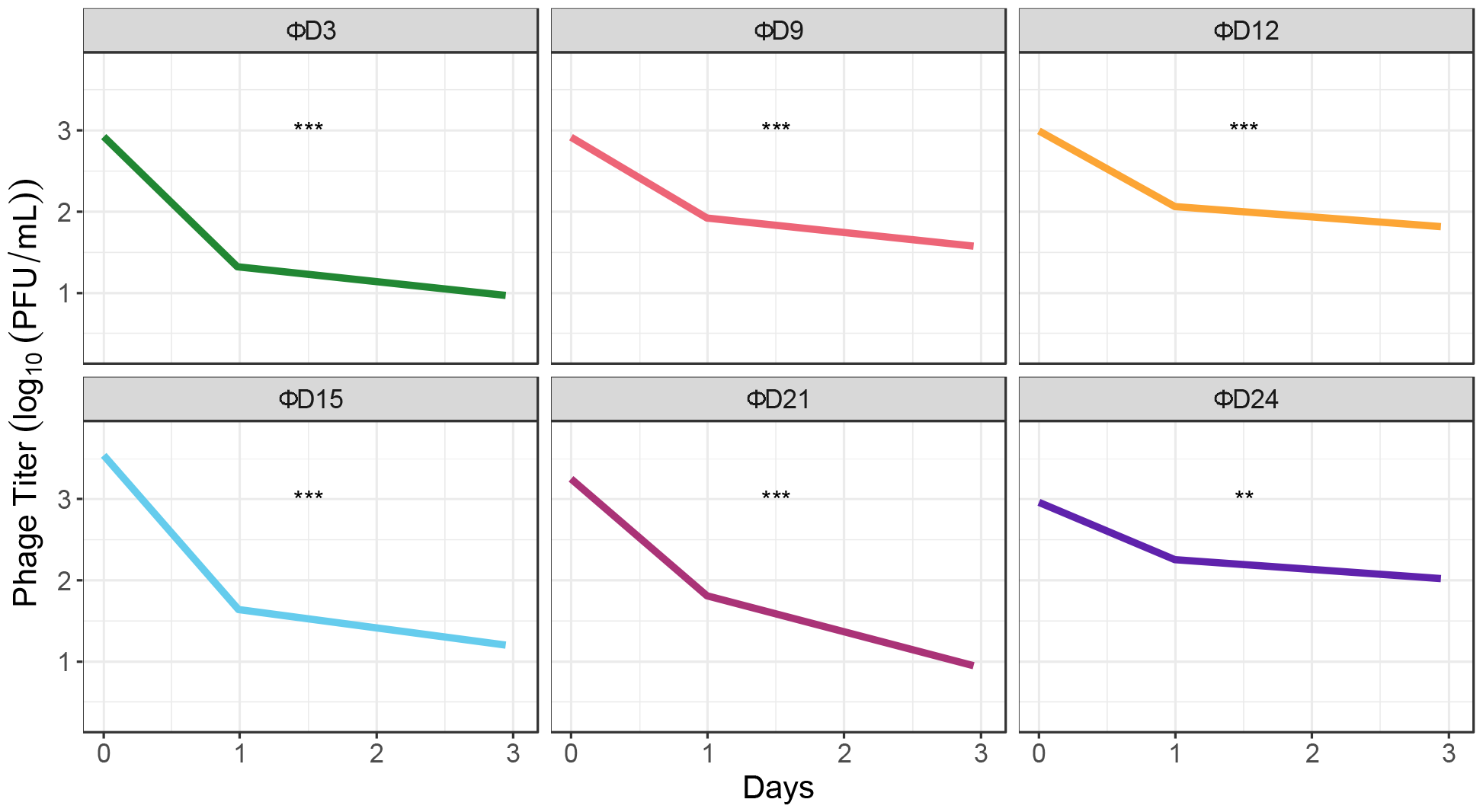
Evolved Φ21 isolates with different host range breadths all demonstrate multiphasic decay. Phage titers were enumerated over 3 days of decay at 37° C. Bold lines represent the mean and faint lines represent independent replicates (n=9). Titer was measured by plating phages and enumerating plaques on soft agar lawns of WT cells (for ΦD24, which only uses the OmpF receptor, we used lawns of ΔLamB ΔOmpC cells). Differences in decay rates between days 0–24 h and 24–72 h are indicated by asterisks (^**^, p<0.01; ^***^, p<0.001; paired T-test).

**Figure 4.**
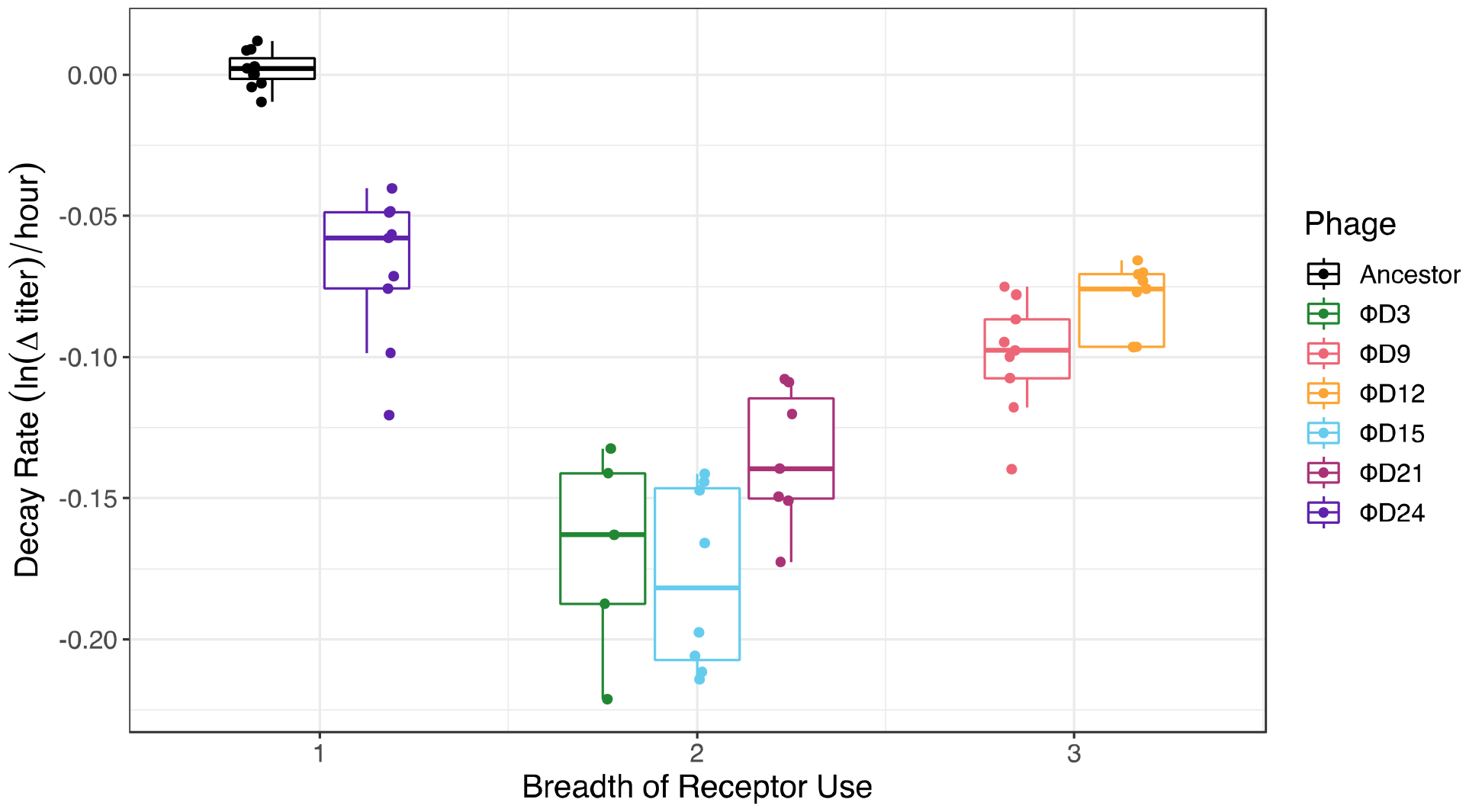
Relationship between breadth of host-receptor use and decay rate of 7 phage genotypes with different receptor-use capabilities. Dual receptor-using phages were the least stable. Receptor-use breadth was linearly correlated with decay rate when including the Φ21 ancestor (linear model; decay rate ~ breadth of receptor use, p=0.001). However, without the ancestor, this relationship is not significant (linear model; decay rate ~ breadth of receptor use, p=0.86).

Next, we explored how the evolution of destabilization relates to different characteristics to better understand the drivers of this key trait. First, we tested whether there was a relationship between the number of receptors a phage can use (receptor breadth) and how unstable it is. We expected that there may be an inverse relationship between thermostability and functional heterogeneity. We found that, by day 3, there is destabilization and the production of phenotypic heterogeneity. Yet, beyond day 3, the relationship between receptor use breadth and stability was not significant (ANOVA, p>0.05). Although Φ21 becomes unstable when it evolves to use the second receptor (OmpC, from ancestor to ΦD3), it then increases in stability when it evolves to use a third receptor (ΦD3 to ΦD9). Then it loses stability again as it evolves in two separate lineages that lose either LamB (ΦD12 to ΦD15) or OmpC (ΦD12 to ΦD21). At later timepoints, Φ21 evolves to specialize on OmpF, becoming the most stable of the evolved phage isolates, yet it is still less stable than the ancestor, which specializes on the LamB receptor.

We also tested whether the number of evolved J mutations or the duration of coevolutionary time were related to stability. In both models, we found that statistical significance was contingent on inclusion of the Φ21 ancestor (Fig. 5A; linear model, decay rate ~ number of J mutations, with Φ21 ancestor p<0.01 & negative slope, without Φ21 ancestor p>0.05 & positive slope) (Fig. 5B; linear model, decay rate ~ days coevolved, with Φ21 ancestor p<0.01 & negative slope, without Φ21 ancestor p<0.05 & positive slope). For the relationships we tested, removing the ancestor from analyses eliminated statistical significance and could alter the direction of the linear relationship. We suspect that the absence of strong relationships between stability and other phenotypic characteristics may be due to the conflicting tension between selection to produce heterogeneity and explore new functions, and costs associated with producing conformers that do not match the availability of receptors on their contemporary evolving hosts. This idea has been demonstrated in influenza RBP (hemagglutinin) evolution, where viruses adapt along a thermodynamic boundary through protein destabilizing and compensatory mutations (Gong et al. 2013).

**Figure 5.**
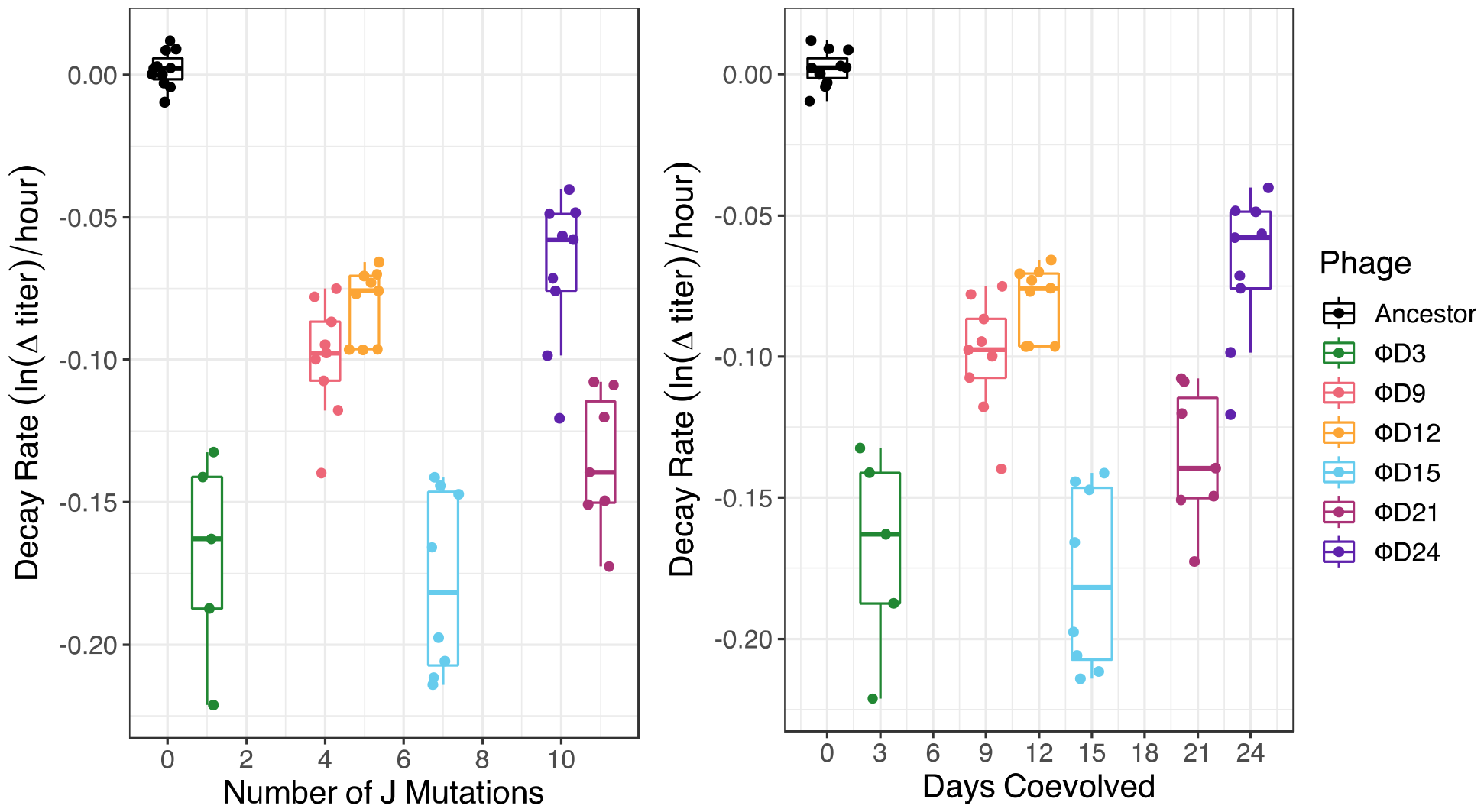
Relationship between phage decay rate and the number of evolved J mutations (Panel A) or days of coevolution (Panel B). Decay rate was calculated as the exponential change in phage titer per hour over the first 24 hours of decay.

### Protein structure predictions

The earliest isolate from the experiment (ΦD3) was sampled after only 3 days of coevolution and had a single mutation in its genome located in the RBP (Table 1). This strain gained the ability to infect using the OmpC receptor, was one of the least stable variants studied, and was bistable, demonstrating that the first evolutionary step taken by the phage simultaneously destabilized the RBP, generated phenotypic heterogeneity, and conferred new host receptor tropisms. This mutation caused a shift from isoleucine to threonine at position 1025, just before the receptor-binding domain is predicted to start. Notably, this mutation is in the same region as destabilizing mutations in λ’s RBP that are essential for gain of function on its new receptor, OmpF (Strobel et al., 2022). Recently, Strobel et al. (2024) demonstrated that, in λ, amino acids in this region of the RBP are involved in trimer formation and substitutions can destabilize hydrogen bonds between trimers, interfere with trimer formation, and cause both instability and gain of function (Strobel et al., 2024). Therefore, we hypothesized that the I1025T mutation may have similar effects on Φ21. We tested our hypothesis by modeling Φ21’s RBP structure and trimer formation. We found that residues I1025 and V1026 are oriented into the center of the tail fiber tube at its distal end, just before the receptor binding domains form (Figure 6). These residues likely stabilize the interaction between the three protomers’ receptor-binding domains and mutating them may alter the stability of these domains’ associations. Given I1025’s precarious location at the junction between domains, weakened interactions from mutation of this residue could allow the receptor-binding domains more conformational freedom to dynamically bind receptors.

**Figure 6.**
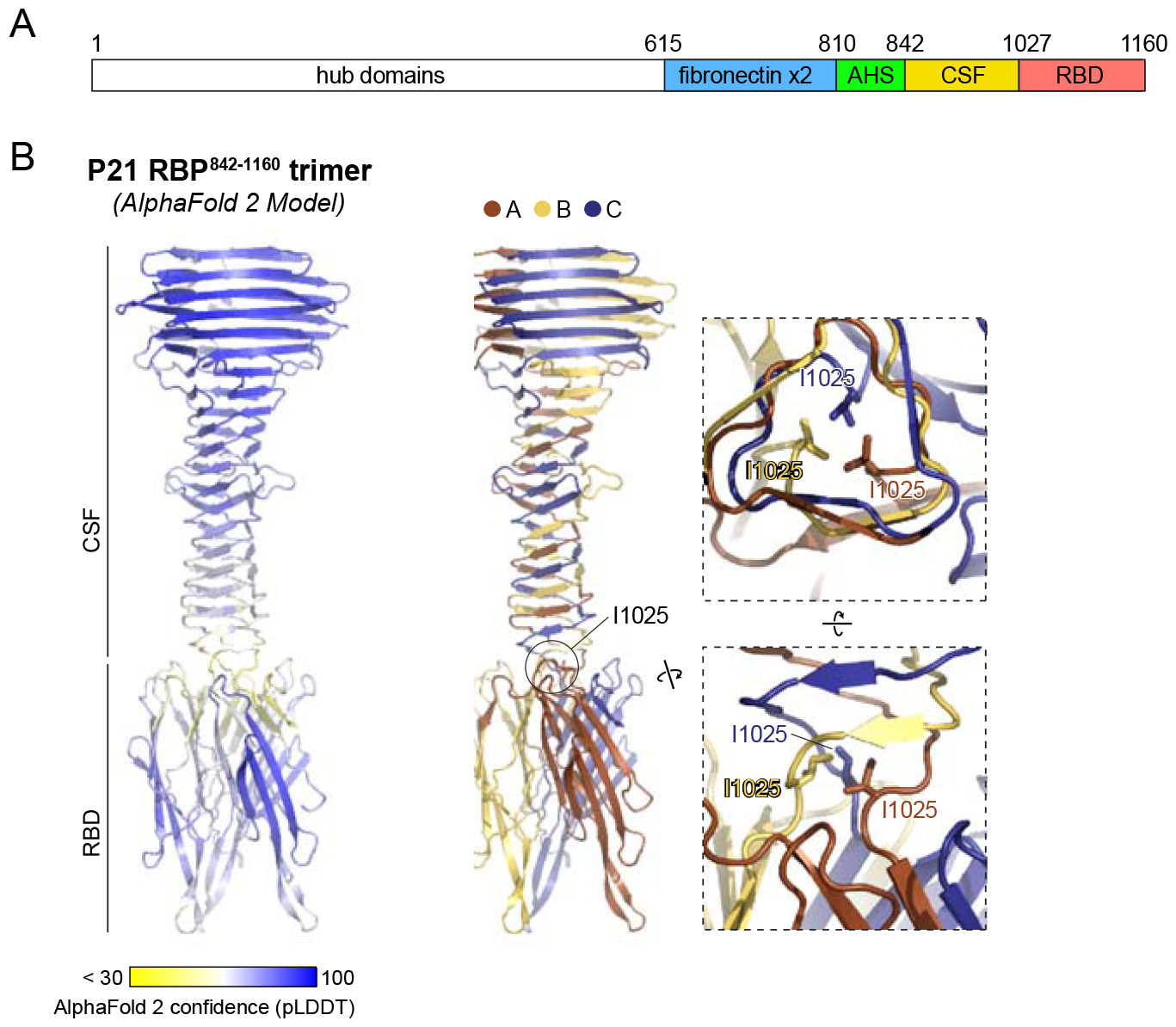
Structure prediction of the Φ21 RBP C-terminal domain trimer. Panel A: Domain schematic of the Φ21 RBP predicted by sequence alignments and structure predictions. Panel B: *Left:* Predicted structure of the Φ21 RBP central straight fiber domain (CSF) and receptor-binding domain (RBD), colored by AlphaFold 2 confidence (pLDDT) scores. *Center:* Predicted structure of the Φ21 RBP CSF and RBD with individual protomers colored brown, yellow, and blue. The position of residue I1025 is highlighted. *Right:* Two closeup views of three I1025 residues making hydrophobic interactions at the central trimer interface of Φ21 RBP.

## Discussion

Viruses often evolve to infect new hosts by modifying their ability to interact with new host receptors (Tétart et al., 1996; Yehl et al., 2019; Boon et al., 2020). Previous work on phage λ demonstrates that viruses can evolve host receptor expansion through the creation of multimorphic intermediate phenotypes that allow the exploration of novel host tropisms. Here we show that a new phage, Φ21, takes a similar path as it evolves to use two new receptors. First, we demonstrate that the receptor binding protein (RBP) of ΦD9, a descendant of Φ21, loses thermostability as it broadens its receptor range and gains the ability to use two new receptors. As in evolved strains of phage λ, the evolved ΦD9 genotype produces a heterogeneous population of phage particles comprised of RBPs with different protein folds that have different thermostabilities and concomitant host receptor tropisms. To test the link between protein folding and host receptor use (function), we manipulated the presence of host chaperone proteins that assist disordered proteins in achieving more stable conformations. These experiments showed that when chaperones were present, phage proteins were more likely to fold into conformations with greater stability that also have a stronger affinity for the native receptor, LamB. Finally, by measuring the decay patterns of 5 additional phage genotypes, with different receptor-use preferences and accumulated RBP mutations, we showed that heterogeneity in phage particles was maintained throughout Φ21’s evolution as its receptor tropism changed to produce a diverse array of receptor-use types. Lastly, we proposed a likely cause of the instability, a mutation that weakens trimer formation at the interface between CSF and RBD domains.

Consistent with previous research conducted on λ, we see that shifts in viral RBP tropism through destabilizing mutations allow for the evolution of new functions while still conferring the ability to infect through the native receptor. Previous studies have referred to the multi-receptor using phage particles as non-genetic phenotypic heterogeneity, and this non-genetic variation has been found to drive evolution (Petrie et al., 2018). Here, we have shown that this process of viral receptor binding protein variation readily evolves in Φ21 as well. Moreover, by altering the heterogeneity of phage receptor-use types by manipulating of host protein-folding chaperones, we have expanded our understanding of how this evolutionary process occurs.

This research also has implications for understanding features that affect protein evolvability. The traditional view is that more stable proteins are more likely to innovate, since they can accommodate more adaptive mutations (mutational robustness) that incur a pleiotropic cost of destabilization (Wang et al., 2002; Bloom and Arnold, 2006; Studer et al., 2014). However, there is a growing realization that the relationship between protein mutational robustness and evolvability may be more nuanced. In some cases, less-stable proteins may be more evolvable if the instability that underlies the formation of phenotypic heterogeneity can be used for functional innovation (Tokuriki and Tawfik, 2009; Sikoset and Chan, 2014; Dellus-Gur et al., 2015). In support of the hypothesis that unstable proteins could enable–rather than hinder–the evolution of new functions, we find that the very first mutation Φ21 evolves destabilizes the phage particle and produces heterogeneous phage particles that allow it to use the novel receptor, OmpC. Even though this observation is not a direct test of this hypothesis, our results support that destabilized proteins can be more evolvable than their structurally rigid counterparts (see Strobel et al., 2022 for a direct test with λ’s RBP). This role of instability promoting protein evolvability does not appear to be limited to viral RBP, as a similar phenomenon has been observed in the evolution of new alkaline phosphatase activity in *E. coli* (Sakuma et al., 2023).

Multiple laboratory studies have demonstrated that RBPs can evolve new receptor tropisms by acquiring destabilizing mutations. This phenomenon could be a product of laboratory evolution. Yet, analyses of natural λ J-related protein sequences show heightened rates of sequence evolution at the same loci that evolve during the laboratory experiments, suggesting similar evolutionary processes are deployed in natural ecosystems (Maddamsetti et al., 2018). Likewise, a systematic analysis of viral proteins suggest that they are generally more disordered and prone to thermodynamic instability than their cellular counterparts (Kramer et al., 2006), which could make phenotypic heterogeneity, and therefore evolutionary innovations, more accessible. Altogether, these observations suggest that destabilization and the production of disordered proteins may be a more common path of viral evolution than has been appreciated.

## Supplementary Materials

### Population decay model selection

To assess whether populations of ancestral phage Φ21 and evolved phage ΦD9 demonstrate monophasic, biphasic, triphasic or continuous decay patterns, we fit 4 models (Eq. 1-4) to each population-level decay curve using the R gslnls package (Figure S1). To differentiate between nested models (Eq. 1-3), we performed log-likelihood ratio tests (LRT). To differentiate between non-nested models, we selected the model with the lowest Akaike Information Criterion (AIC). For the Φ21 ancestor, Model 1 (e.g., monophasic decay, Eq. 1) was selected as the best fit model. The addition of parameters for bi-and tri-phasic decay did not significantly increase the variance explained in the population level decay (P>0.05, LRT). Further, the AIC of Model 1 is lower than that of Model 4, confirming the choice of monophasic decay as the best fit of these models to this data (Table S1). For ΦD9, Model 2 (e.g., biphasic decay, Eq. 2) was selected as the best fit model. Eq. 2 significantly improves the variance in population decay explained relative to Eq 1. (p<0.01, LRT); however, Eq. 3 does not significantly increase the goodness of fit over Eq. 2 (p=0.78, LRT). Additionally, the AIC of Model 2 is lower than that of Model 4, further supporting that ΦD9 demonstrates biphasic decay.

---

**Figure S1.**
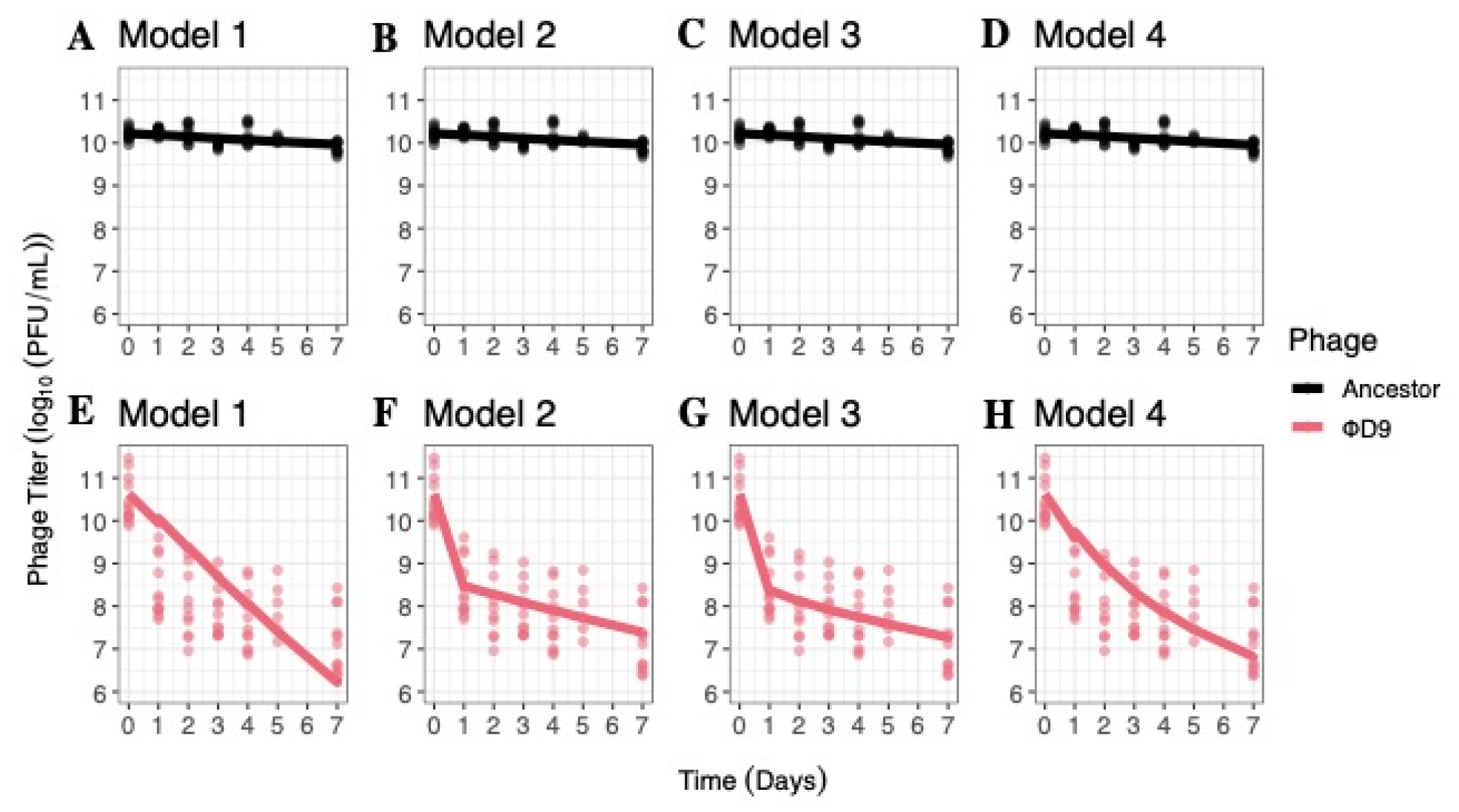
Exponential decay models indicate monophasic decay curves in the Φ21 ancestor and biphasic decay curves in the evolved ΦD9 phage. Models 1–4 were fitted based on equations 1–4 in the main text and represent one, two, or three population decay rates, or a continuously decaying model, respectively. Panels A–D: Models 1–4 fit to Φ21 ancestor decay over 7 days. Panels E–H: Models 1–4 fit to ΦD9 decay over 7 days. Model 1 was selected as the best fit to the decay pattern of the Φ21 ancestor over 7 days. Model 2, indicating biphasic decay, was the best fit model to the decay pattern of the evolved ΦD9 phage.

**Table S1.** Φ21 ancestor and ΦD9 decay patterns analyzed with Akaike Information Criterion (AIC) for model selection. Table denotes phage ID, model number, and its relevant AIC. The lowest AIC, determined through fit and model complexity, of each phage ID suggests the most supported model.

| Phage | Model Fitted | Akaike Information Criterion (AIC) |
| --- | --- | --- |
| Φ21 ancestor | Model 1 | -39.5649 |
| Φ21 ancestor | Model 2 | -33.5649 |
| Φ21 ancestor | Model 3 | -29.5649 |
| Φ21 ancestor | Model 4 | -38.52257 |
| ΦD9 | Model 1 | 228.5354 |
| ΦD9 | Model 2 | 153.2392 |
| ΦD9 | Model 3 | 157.7206 |
| ΦD9 | Model 4 | 198.7129 |

## Acknowledgements

We acknowledge Katherine Petrie and Lin Chao for helpful input, Sweetzel Labador for laboratory assistance, Hannah Strobel for decay protocols, the National BioResource Project (NIG, Japan) for the ASKA strain, and funding through Howard Hughes Medical Institute Emerging Pathogens Initiative.

*Data available at https://doi.org/10.5061/dryad.wwpzgmsrw*

